# Heat denaturation enables multicolor X10-STED microscopy at single-digit nanometer resolution

**DOI:** 10.1101/2022.06.24.497479

**Authors:** K.-A. Saal, A.H. Shaib, N. Mougios, D. Crzan, F. Opazo, S.O. Rizzoli

## Abstract

Expansion microscopy (ExM) improves imaging quality by physically enlarging the biological specimens. In principle, combining a large expansion factor with optical super-resolution should provide extremely high imaging precision. However, large expansion factors imply that the expanded specimens are dim and are therefore poorly suited for optical super-resolution. To solve this problem, we present a protocol that ensures the 10-fold expansion of the samples through high-temperature homogenization (X10ht). The resulting gels exhibited relatively high fluorescence intensity, enabling the sample analysis by multicolor stimulated emission depletion (STED) microscopy, for a final resolution of 6-8 nm. X10ht offers a more thorough homogenization than previous X10 protocols based on enzymatic digestion, and thereby enables the expansion of thick samples. The better epitope preservation also enables the use of nanobodies as labeling probes and the implementation of post-expansion signal amplification. We conclude that X10ht is a promising tool for nanoscale resolution in biological samples.

## Introduction

Imaging cellular proteins at a resolution similar to their sizes, around 1 to 10 nm, is still a challenge, in spite of two decades of progress in super-resolution optical microscopy (see for example reviews [1,2]). This type of resolution is only achieved regularly by highly specialized optics approaches as MINFLUX imaging (maximally informative luminescence excitation, [3–5]), which are not yet widely available. In principle, the same type of resolution should be achievable by taking advantage of an unconventional super-resolution method, expansion microscopy (ExM), which has been introduced by Boyden and collaborators [6]. In ExM, a specimen is embedded in a swellable gel, is homogenized in a procedure that severs the connections between its components (and/or severs the proteins themselves), and is then expanded in an isotropic fashion. The enlargement of the biological sample via ExM circumvents the diffraction limit of light, by physically enlarging the distances between the fluorophores decorating the targets (reviewed by [7]).

The resolution obtained with ExM approaches scales inversely with the expansion factor, starting around 70 nm in the initial implementations of 4-fold ExM [6] and reaching around 25 nm with 10-fold implementations [8–10]. The resolution is enhanced further by iterative procedures that expand cells 10-20 fold (iExM, [11]), albeit the repeated expansion procedures may result in distorted specimens, which ultimately affects not only resolution, but also target identification. Overall, the resolution obtained when relying exclusively on ExM procedures peaks around 20 nm, rarely going beyond this value.

In principle, a simple procedure for further enhancing resolution would be the combination of ExM with conventional super-resolution techniques. This has been attempted many times, relying typically on the original 4× protocol of the Boyden laboratory, using, for example, stimulated emission depletion microscopy (STED) [12–14], stochastic optical reconstruction microscopy (STORM) [15–17], or structured illumination microscopy (SIM) [18–20]. However, the resulting imaging precision is often only slightly higher than that obtained in optimal implementations of STORM microscopy [21,22]. While some of the ExM-optical super-resolution protocols reached resolutions of 4-8 nm in individual imaging channels, using either STED or STORM (*e.g*. [12,17]), the use of 4× ExM seems to limit imaging precision, requiring the use of gels with larger expansion factors.

Such gels have been more difficult to combine with optical super-resolution, since large expansion factors result in dim samples that are difficult to image with the challenging optics used for STED, STORM or SIM, which is simply due to the low fluorophore densities [23,24]. Moreover, large expansion factors require a thorough homogenization of the samples, relying on enzymatic digestion with molecules as proteinase K, which results in a loss of proteins and fluorophores from the samples, further decreasing the sample intensity. To address this issue, we combined here a 10-fold expansion protocol (X10, [8,25]) with sample homogenization by heating in alkaline conditions, as introduced for 4× ExM by Boyden and collaborators [26]. This protocol, which we termed X10 heat-treated (X10ht), enables a much better preservation of fluorophores in the samples, and therefore allows the use of the samples in multicolor STED imaging, resulting in resolutions below 10 nm in at least three color channels simultaneously. At the same time, X10ht deals much better with thick tissue specimens than the original X10 protocol, implying that it should be a more promising imaging technique for such samples.

## Results

### 2.1 X10ht microscopy

X10 microscopy enables a resolution of ~20-25 nm [8], but was somewhat limiting in the choice of the samples. The gel polymers based on *N,N*-dimethylacrylamide acid (DMAA) and crosslinked with sodium acrylate (SA) form a dense matrix in thick specimens, which prevents the penetration of proteases, and limited X10 to very thin tissue slices, up to ~10 μm, or to cell cultures [8]. To address this problem, we turned to treating the samples at high temperature in a detergent-rich buffer, at alkaline pH [26]. Such conditions unfold proteins, disengaging protein-protein interactions, and also break peptide chains by hydrolysis [27], resulting in a thorough sample homogenization.

To avoid the loss of proteins and fluorophores from the gels, we first maximized the anchoring of lysine residues to the gel matrix by using the succinimidyl ester of 6-((acryloyl)amino))hexanoic acid (acryloyl-X, AcX). This molecule, which conjugates itself to the free (exposed) amines of lysine residues and of protein N-termini, is employed in most ExM publications, and was applied here at a 3-fold higher concentration than in the original X10 protocol (0.3 mg/ml), in a buffer designed to enhance its reactivity to the maximum possible (strongly alkaline conditions). Heat treatments were then applied using an autoclave, which enabled the analysis of different temperatures, without evaporation of the sample buffers. Heating at 70°C or above resulted in relatively strong cellular expansion, but the cells exhibited cracks and/or swelled regions (“bubbles”), until the heat treatment reached at least 100-110°C (Supp. Fig. S1). Importantly, homogenization via autoclaving at 110°C led to a similar expansion factor to the original X10 protocol (based on proteinase K digestion), as shown in Supp. Fig. S2.

We then investigated the performance of the two protocols in fluorescence imaging. We immunostained microtubules, relying on a protocol combining several primary antibodies (described in Methods), and compared expanded samples after subjecting them to X10 or X10ht. For simplicity, in the figures we use refer to the two homogenization methods as “autoclave treatment” (AC) or “proteinase K treatment” (PK). The heating-based protocol resulted in substantially brighter samples, which were much easier to image in both confocal and STED microscopy (Fig. 1), implying that X10ht limits the loss of fluorophores through excessive homogenization.

**Figure 1.**
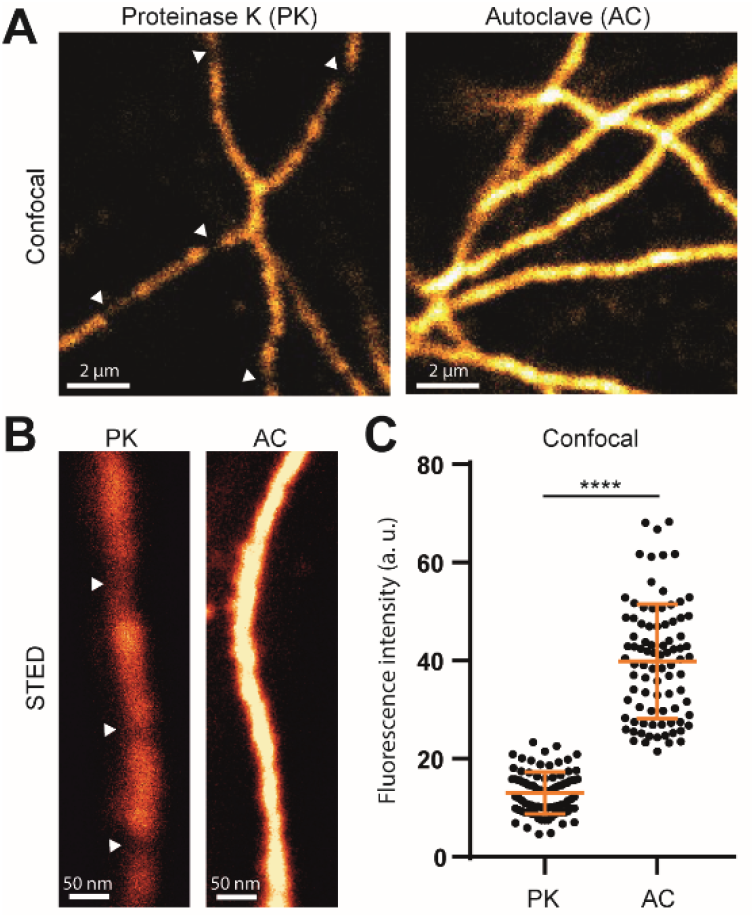
X10ht enables confocal and STED imaging of tubulin immunostained in U2OS cells. **A)** Representative images of tubulin under confocal or **B)** STED imaging, indicating the difference in fluorescence intensity in both methods (autoclave treatments, AC, and proteinase K treatments, PK). Using AC, we observed that the fluorescence signal appeared as a continuous decoration of the microtubules, while with PK the loss of fluorophores results in “broken” microtubules (white arrows). N = ~90 regions of interest (ROIs). Data are presented as single ROI data points, mean ± SD.

We therefore conclude that heat treatments are suitable for X10 gels, resulting in a functional X10ht expansion protocol.

### 2.2 X10ht enables usage of nanobodies as labeling probes

Single domain antibodies (nanobodies, [28]) are substantially smaller than antibodies, thereby placing the fluorophores much closer to their targets (e.g. [29]), which makes them preferable to antibodies in many immunostaining approaches [30,31]. However, the small size of these tools renders them difficult to link to expansion gels [12], since they contain only a handful of exposed amines. To test the usability of nanobodies in the X10ht protocol, we started by immunostaining primary neuronal cultures with primary antibodies that were detected by secondary, fluorophore-conjugated nanobodies. Digestion with proteinase K resulted in very weak labeling of the neuronal structures (Fig. 2A), but this increased strongly under X10ht conditions (Fig. 2A,B). We then targeted the protein of interest (POI) directly with nanobodies labeled with AlexaFluor 488 (AF488). Immunostainings for the vesicular glutamate transporter1 (VGLUT1) were invisible after proteinase K homogenization, but could be easily detected in X10ht (Fig. 2C,D), indicating that X10ht should be the preferred protocol for such labeling approaches.

**Figure 2:**
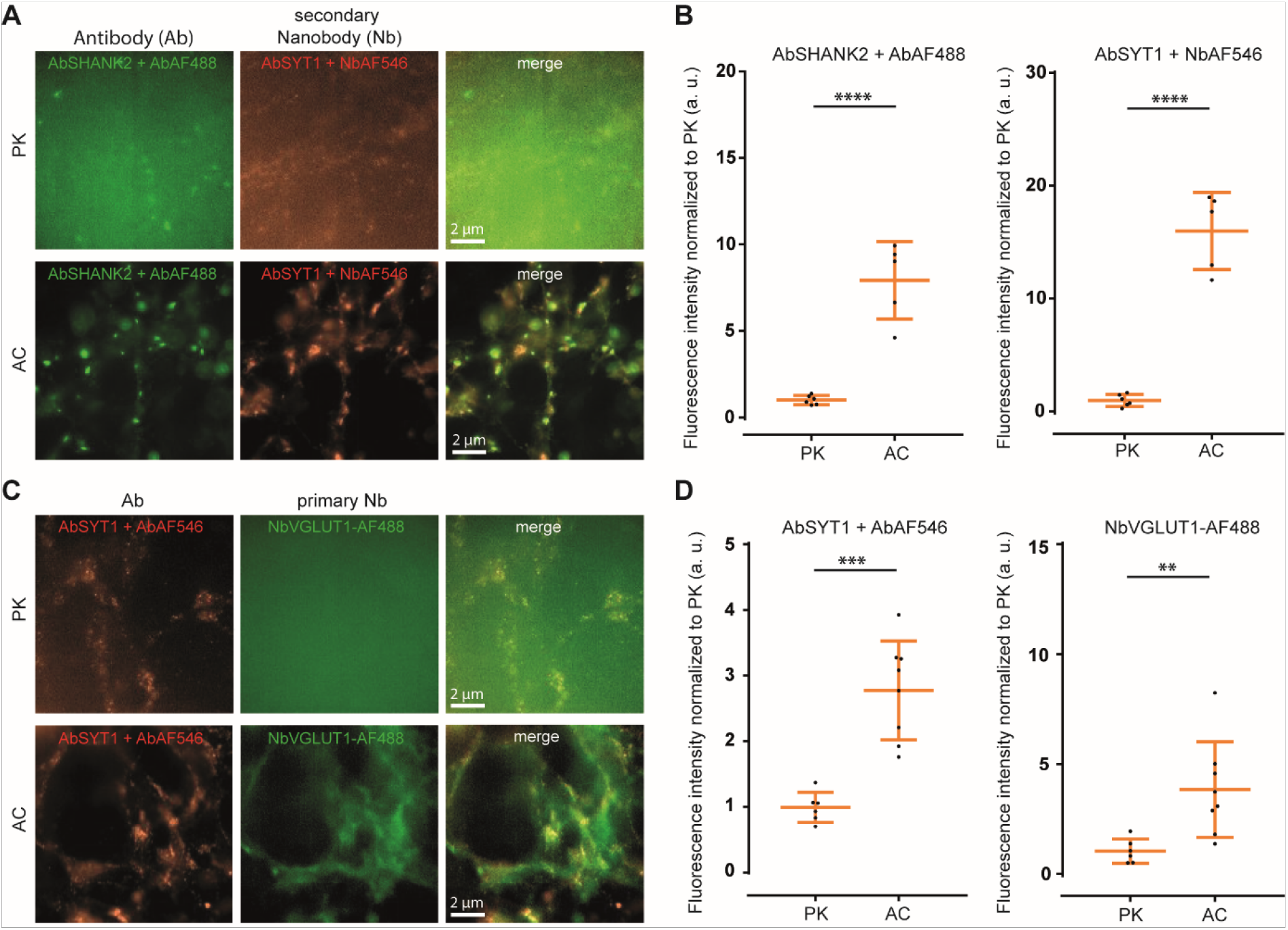
Applicability of nanobodies in X10ht, compared to homogenization with proteinase K (original X10 protocol). **A)** Exemplary epifluorescence images of hippocampal cultured neurons immunostained with antibodies (Ab) for the postsynaptic protein SHANK2 and the presynaptic protein synaptotagmin1 (SYT1), revealed by secondary antibodies (AbSHANK2 + AbAF488) or secondary nanobodies (Nb) (AbSYT1 + NbAF546). **B)** A pronounced increase in fluorescence intensity in both immunostainings is obtained when samples are autoclaved (AC) compared with the original X10 protocol (PK). N = 6 images with several ROI analyzed for AbSHANK2 + AF488 PK and AbSYT1 + NbAF546 PK, and N = 5 images with several ROI analyzed for AbSHANK2 + AF488 AC and AbSYT1 + NbAF546 AC. **C)** Exemplary images of SYT1 antibody immunostainings, revealed by a secondary antibody (AbSYT1 + Ab AF546), in parallel with labeling for the vesicular glutamate transporter 1 (VGLUT1) with primary nanobodies coupled to AF488 (NbVGLUT1-AF488). **D)** The quantification of the fluorescence signal revealed significantly increased intensities with X10ht. N = 6 images with several ROI analyzed for AbSYT1 + AbAF546 PK and NbVGLUT1-AF488 PK, and 8 images with several ROI analyzed for AbSYT1 + AbAF546 AC and NbVGLUT1-AF488 AC. Data are presented as single data points, mean ± SD.

### 2.3 X10ht allows the analysis of thick tissues

Since the original X10 protocol did not allow the homogeneous expansion of thick tissue slices, we tested X10ht on fixed rat brains, using 100-200 μm slices. We immunostained the samples using antibodies detecting the pre- and postsynaptic compartments, followed by the X10ht procedure (Fig. 3). As expected, homogenization with proteinase K failed to facilitate isotropic expansion of the medial tissue area, resulting in the gels disintegrating to some extent during the expansion procedure [8,25]. X10ht led to the macroscopic preservation of the tissue organization (Fig. 3A,C), albeit the tissues typically reached only ~6-fold expansion (Fig. 3G,H), failing to reach the full potential of the expansion gel. This lower expansion factor in thicker specimens is a known ExM phenomenon when working with large tissue samples, and reflects difficulties in homogenizing the thick connective tissue bundles.

**Figure 3:**
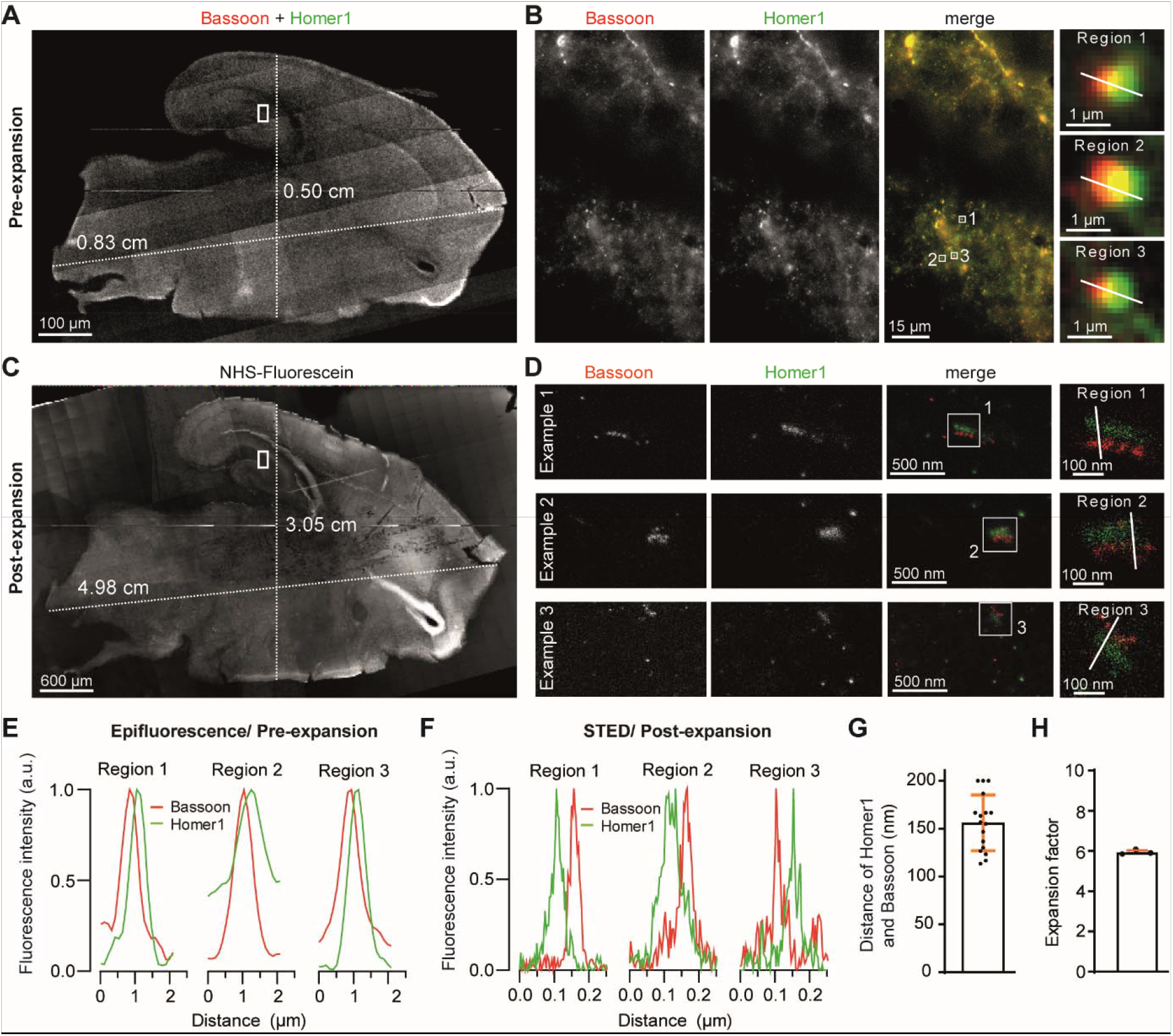
X10ht of 200 μm thick rat brain slices immunostained for the synaptic proteins Bassoon and Homer. **A)** Epifluorescence tile image of merged channels displays a non-expanded 200 μm thick rat brain slice, containing the region of the hippocampal formation. **B)** The region marked (white rectangle) in panel A is shown here, indicating the immunostaining for Bassoon (red) and Homer1 (green). The marked regions (1-3) are shown in the insets. **E)** Plots of the line scans over the zoomed areas. **C)** Epifluorescence tile acquisition of the same brain slice after expansion labeled with NHS-Fluorescein, depicting the retention of tissue shape and the prolongation of the slice length from 0.83 to 4.98 cm and width from 0.5 to 3.05 cm. **D)** Representative STED images of different regions from the C1 region of the hippocampus, showing highly resolved pre- (red) and postsynaptic (green) compartments. **F)** The respective line scans are plotted, and the distance between the pre- and postsynaptic compartments is analyzed in **G)**. N = 17 synapses. **H**) The graph shows an expansion factor of 6 with N = 3 gels of two independent experiments.

Nevertheless, the good preservation of the immunostaining in X10ht implied that the samples could then be subjected to 2-color STED imaging, resulting in excellent resolution. The pre- and postsynaptic compartments are easily observed in the resulting images and are clearly separated by the expected distance (Fig. 3D,F,G; [8,25]).

### 2.4 Amplifying the signal intensity with different amplification systems

Albeit the samples treated by X10ht can be used in STED, they are still substantially dimmer than non-expended samples, due to the 1000-fold (for fully expanded samples) dilution of the fluorophores in the expanded volume. To amplify signal intensities, we devised several methods, which we combined to perform multicolor ExM (Fig. 4).

**Figure 4:**
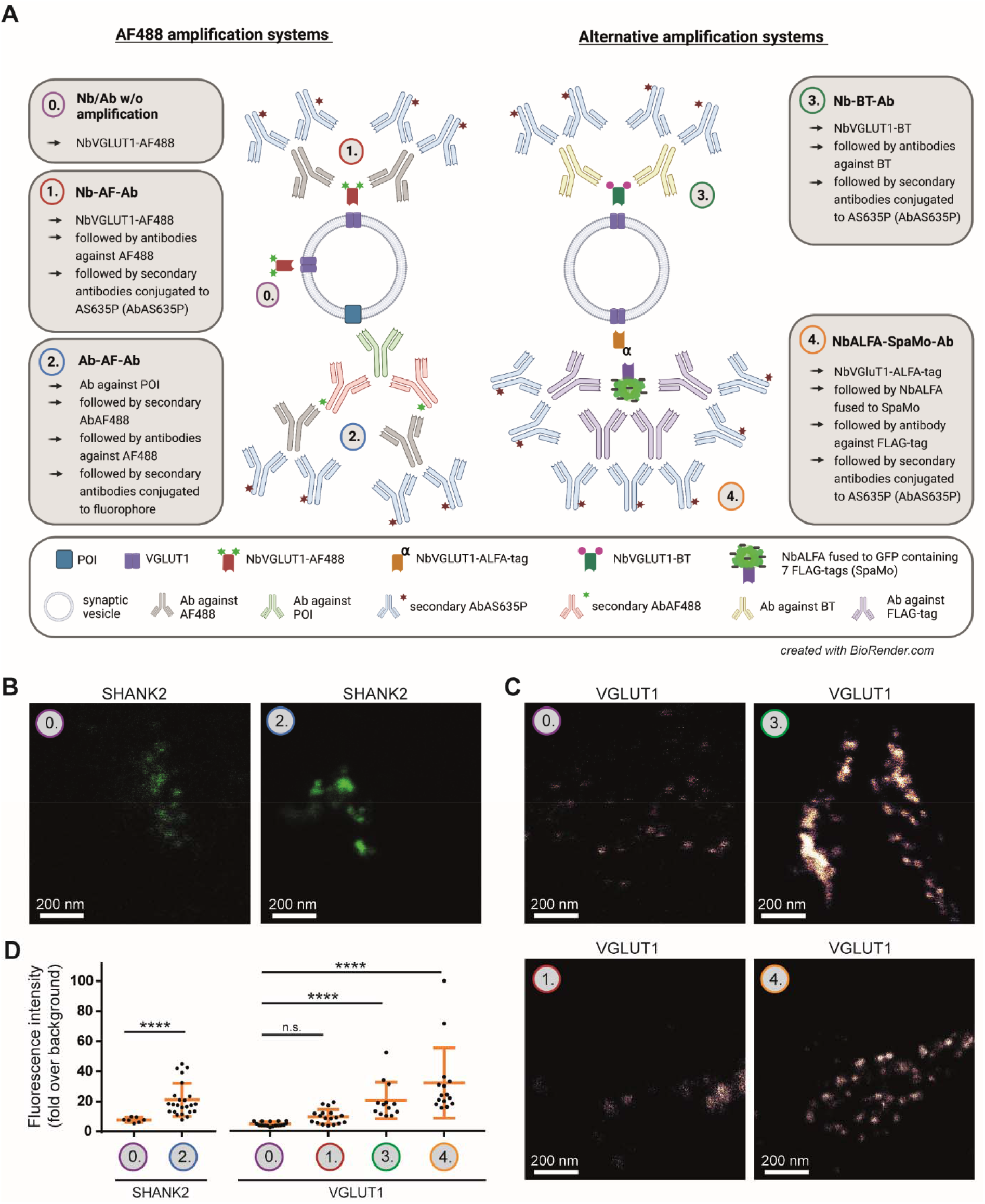
Pre-expansion labeling of presynaptic markers with different signal amplification systems in X10ht. **A)** Schematic visualization of the different amplification systems. See the main text for more details. Final used fluorophores are Alexa Fluor488 (AF488) or Abberior Star635P (AS635P). **B)** Representative confocal images of expanded neurons immunostained with either the antibody-AF488-based amplification system for SHANK2, or **C)** with the different nanobody-based amplification systems detecting VGLUT1. **D)** Quantitative analysis of SHANK2 with N = 7 images for 0. Ab w/o amplification, and N = 22 images for 2. Ab-AF-Ab; N = 18 images for 0. Nb w/o amplification (VGLUT1) and 1. Nb-AF-Ab, N = 13 images for 3. Nb-BT-Ab, and N = 14 images for 4. NbALFA-SpaMo-Ab all to detect VGLUT1, with several ROIs analyzed per image. Data are presented as single data points, mean ± SD.

We first used Alexa Fluor 488 (AF488) as an amplification tool. Samples were immunostained pre-expansion with a primary nanobody conjugated to AF488 (NbVGLUT1-AF488), which was then detected by an antibody against AF488, which was in turn revealed by secondary antibodies carrying a second fluorophore (Fig. 4A, protocol 1., “Nb-AF-Ab”). Another approach used an unconjugated primary antibody, detected by AF488-conjugated secondary antibodies, which were in turn revealed by antibodies against AF488 and their specific secondary antibodies (Fig. 4A, protocol 2., “Ab-AF-Ab”). A third approach focused again on primary nanobodies, replacing AF488 with biotin (BT) (Fig. 4A, protocol 3., “Nb-BT-Ab”). Finally, we used primary nanobodies carrying the ALFA-tag [32], which were detected by an anti-ALFA nanobody (NbALFA) fused to a FLAG-tag spaghetti monster, SpaMo [33], an engineered GFP containing seven FLAG-tags. These tags were detected by antibodies against FLAG-tag, followed by secondary antibodies (Fig. 4A, protocol 4., “NbALFA-SpaMo-Ab”).

The protocols based on primary and secondary antibody usage with AF488 amplification resulted in a 2.5-fold signal increase (Fig. 4B,D), while the lowest signal amplification was observed when applying the AF488 system for the detection of VGLUT1 with the primary nanobody (Fig. 4C,D). The BT-based protocol amplified signals ~5-fold, when compared to direct nanobody immunostainings, and the NbALFA-SpaMo-based protocol was the most successful one, resulting in a 7-fold improvement of sample intensity (Fig. 4C,D). In principle, applying these tools after the denaturation process of the gels would further increase signals, by eliminating steric hindrance. We performed this analysis with the NbALFA-SpaMo-based protocol (Fig. 5A), and observed a highly significant increase in fluorescence intensities (Fig. 5B,C), which implies that this approach, based on pre-expansion labeling with ALFA-tag carrying nanobodies, followed by SpaMo-based detection after expansion, is feasible for the analysis of target proteins.

**Figure 5:**
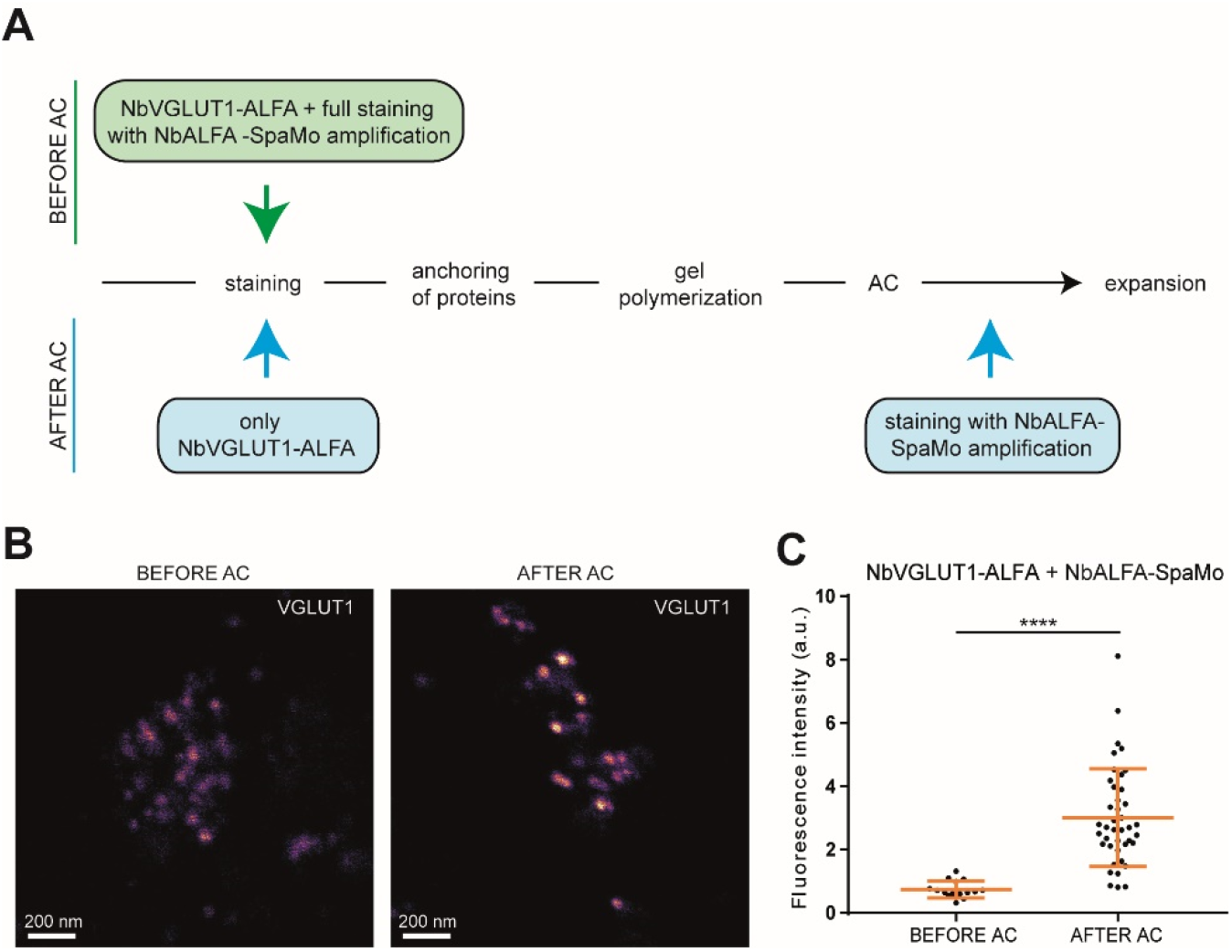
Application of the NbALFA-SpaMo system before and after sample expansion. **A)** Scheme of two possible workflows for the NbALFA-SpaMo application. **B)** Representative confocal images for both NbALFA-SpaMo applications before or after autoclaving (AC). **C)** A quantification of the signal intensities in the different protocols. N = 15 images for before AC and 41 images for after AC with several ROI analyzed, obtained from 2 independent experiments. Data are presented as individual data points, mean ± SD.

The higher fluorescence intensity provided by the signal amplification procedures was then employed to confirm the 10x expansion of the specimens, relying on a well-known structure, the nuclear pore complex (NPC;[24]). We employed a genetically modified knock-in U2OS cell line that expresses mEGFP coupled to the nucleoporin Nup96 [34]. We detected GFP using AF488-conjugated primary nanobodies, whose signal was amplified by primary (anti-AF488) and secondary antibodies. STED images revealed only five to six of the eight NPC subunits, even before expansion (Supp. Fig. S3A,B), most probably due to incomplete labeling [24,34]. To avoid difficulties due to this issue, we focused on analyzing the distance between neighboring Nup96 subunits, rather than on the nuclear pore sizes. As expected from literature [35], they appeared to be ~38 nm apart (Supp. Fig. 3C).

### 2.5 X10ht enables multicolor X10 STED acquisition of synaptic vesicles

The different signal amplification techniques introduced above enabled the multicolor labeling of at least three different targets. Relying on primary neuronal cultures, we used the AF488 amplification system to detect the postsynaptic protein SHANK2 and combined it with labeling of biotinylated primary nanobodies and with the NbALFA-SpaMo system for the presynaptic markers SYT1 and VGLUT1, respectively (Supp. Fig. S4). While the AF488- and the BT-amplification was applied before autoclaving the samples, the SpaMo (along with the respective anti-Flag antibodies) was added after the heat treatment. The pre- and postsynaptic compartments were revealed by this approach, as indicated in Supp. Fig. 4.

To test the resolution of the resulting immunostainings, we turned to a simpler biological sample, isolated synaptic vesicles applied on coverslips, as sparse monolayers (as we used for imaging purposes in the past, e.g. [36]). We combined the amplification systems indicated above for three synaptic vesicle markers, synaptophysin 1 (SYP1), SYT1 and VGLUT1, and could observe the expected patterns for these immunostainings (Fig. 6A,C). Each of the proteins resulted in multiple “spots” on individual vesicles, which correspond to the rough organization of the targets, taking into account the size of the antibody packages (Supp. Fig. S5). The analysis of the fluorescence intensities revealed that all signals were sufficient to enable STED imaging (Fig 6B,D). To obtain an indication of the observable resolution, we measured the size of small spots (presumably individual fluorescently-labeled Fc fragments of secondary antibodies). The values averaged to ~7-8 nm (Fig. 6F), which should be equivalent to the resolution of the X10ht-STED approach.

**Figure 6:**
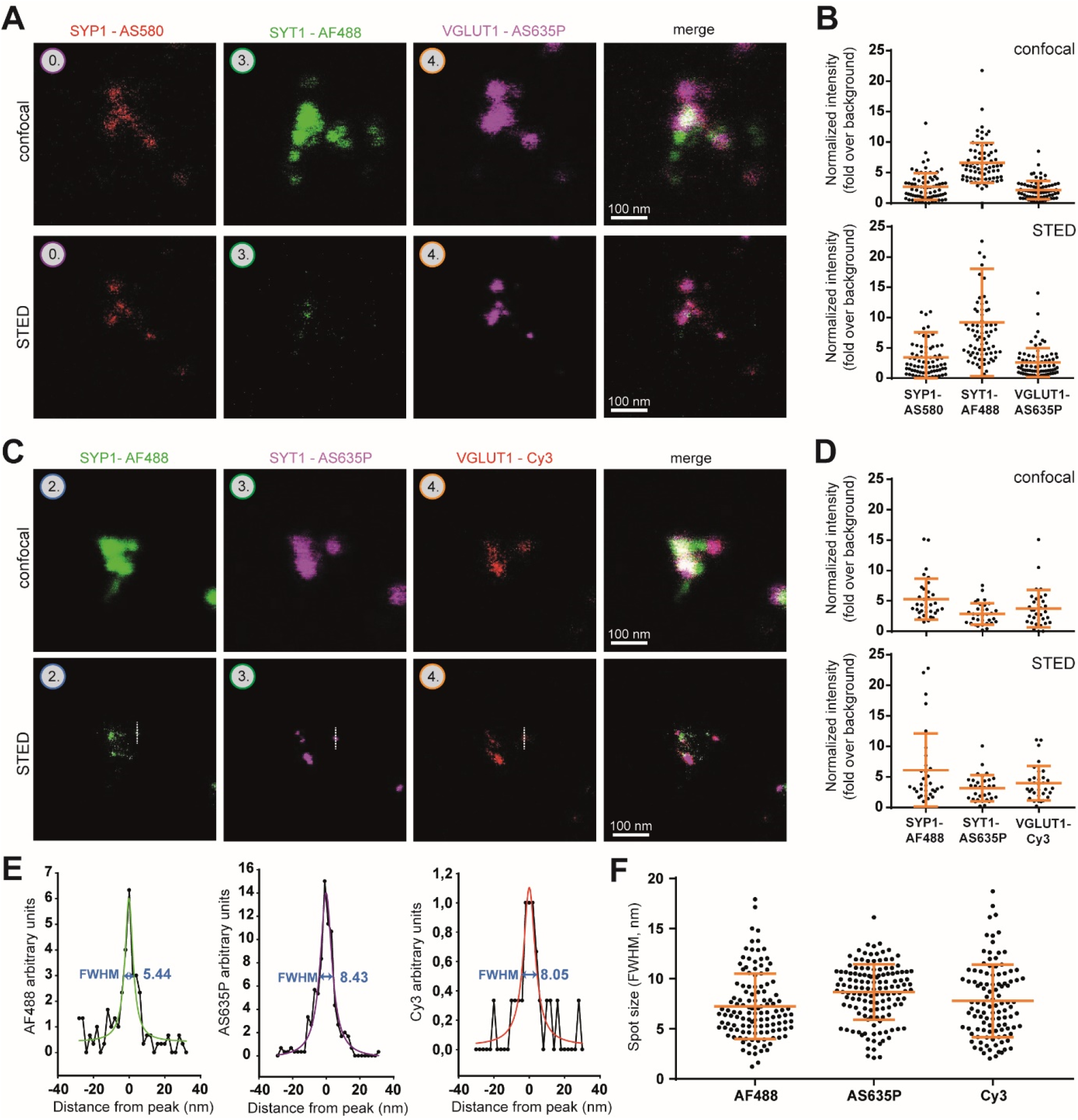
X10ht-STED imaging of isolated synaptic vesicles. **A)** Representative confocal and STED images of one possible combination of amplification systems for SYP1, SYT1 and VGLUT1, revealed by AS580, AF488 and AS635P, respectively. **C)** The protein targets and their amplification systems are combined with another selection of fluorophores. **B), D)** The fluorescence intensities of all channels were analyzed, and indicated a sufficient signal-to-noise ratio for vesicle identification. N = 74 ROIs from 27 images in B, and N = 33 ROIs from 20 images in D. **E)** Line scans were drawn over the spots indicated in C). The respective full widths at half maximum (FWHM) are indicated. **F)** An analysis of spot sizes. N = 72 analyzed AF488 spots from 20 images, N = 78 AS635P spots from 18 images, and N = 34 Cy3 spots from 14 images. Data for B, D and F are presented as individual data points, mean ± SD.

## Discussion and Conclusion

ExM was introduced to offer a facile and widely-applicable option for super-resolution investigations [6], and it has largely fulfilled its potential, with numerous groups adopting it as a technique of choice for biological investigations. At the same time, while ExM applications reach easily 50-70 nm resolutions, higher values require more complex protocols that have been less widely applicable, as iterative ExM (iExM), X10 or TREx [8,10,11]. Even with these expansion protocols, which bring ExM performance to ~20 nm, the domain of single-digit nanometer resolution remains largely untouched, requiring a combination of ExM and advanced optical methods.

This has been often attempted, as mentioned in the Introduction, and has occasionally resulted in excellent resolution (4-8 nm), even when relying on 4x ExM and applying highly advanced microscopy (e.g. [12,17]) provide similar resolutions using any commercial super-resolution setup, one should rely on larger expansion factors. We performed this here, optimizing the original X10 expansion protocol [8,25], changing the homogenization method from proteinase K treatments to autoclave heating in an alkaline milieu. The less aggressive protein fragmentation reduces the loss of fluorophores during the expansion process, maintains small molecules like nanobodies in the gel, and therefore enables more effective imaging in super-resolution, for multiple channels.

To overcome low signals, one could also rely on post-homogenization labeling or signal amplification [37]. Faced with harsh homogenization procedures, it is most efficient to rely on multifunctional anchors [17,38], which improve the fluorophore coupling to the gels structure to levels that are sufficient for optimal ExM-super-resolution combinations [17]. However, a mild homogenization is still preferable, since it retains not only a variety of fluorophores and epitopes, but also the general protein makeup of the cells, which can be labeled fluorescently (e.g. [39]), or the lipid membranes, which have been recently visualized in TREx [10], as well as in sphingolipid ExM [40]. Mild homogenization procedures also enable efficient post-expansion immunostainings, which result in higher intensity levels, since more antibodies fit into the same region of the expanded cell [20,26], to such a level that new structures become visible [41,42].

A surprising effect of the milder homogenization produced by alkaline heating was that it enabled a more thorough expansion of thick specimens (200 μm, versus 5-10 μm in the original X10 protocol). This can be attributed to difficulties in enzyme penetration into thick, gel-embedded specimens, something that is not an issue for heating-based protocols (as already exploited, for example, in ZOOM ExM, [9]).

The nominal resolution obtained by X10ht-STED is suitable for investigating small structures as protein complexes. This type of resolution rivals that obtained with several highly advanced procedures, both in the ExM and the advanced optics fields. However, a caveat is that the use of amplification systems induces a localization error that depends on the type and number of tools used (nanobodies, antibodies), and therefore the effective, usable resolution will depend on the type of labeling employed.

We conclude that the optimized version of the X10 ExM protocol that we established here is a promising approach for super-resolution investigations, and that it is a relatively easy avenue in the direction of multi-color single-digit nanometer investigations.

## Acknowledgements

We thank Christina Zeising and Nicole Hartelt for excellent technical assistance.

## Materials and Methods

### U2OS cell line

This knock-in cell line, which expresses the nuclear pore complex 96 (NUP96) tagged with mEGFP (U-2 OS-CRISPR-NUP96-mEGFP clone #195), was purchased from the CLS cell lines service (Eppelheim, Germany). Cells were maintained at 37°C, 5% CO2 in a humified incubator in Dulbecco’s Modified Eagle Medium (DMEM #D5671, Merck, Darmstadt, Germany) supplemented with 10% Fetal Calf Serum (FCS, #S0615, Merck), 4 mM L-glutamine (#25030-024, ThermoFisher Scientific, Waltham, USA) and 1% penicillin/streptomycin (ThermoFisher Scientific), and split twice a week. For experimental usage, cells were plated on poly-L-lysine (PLL, #P2658, Merck) -coated 18 mm glass coverslips and allowed to grow for 24h.

### BHK cells

Baby hamster kidney (BHK) fibroblasts were cultured in DMEM, containing 10% tryptose phosphate broth solution (#T8159, Sigma-Aldrich now Merck), 5% FCS, 2 mM L-glutamine, 60 U/ml penicillin and 60 U/ml streptomycin. Cells were split twice a week, for experiments seeded on PLL coated coverslips and fixed after ~24 h with 4% PFA in PBS for 30 minutes.

### Primary neuronal culture

Primary hippocampal neurons were obtained from newborn Wistar rats (P0-P1), bred in the animal facility of the University medicine Göttingen. All animals were handled ac-cording to the specifications of the University of Göttingen and of the local authority, the State of Lower Saxony (Landesamt für Verbraucherschutz, LAVES, Braunschweig, Germany). The local authority, the Lower Saxony State Office for Consumer Protection and Food Safety (Niedersächsisches Landesamt für Verbraucherschutz und Lebensmittelsicherheit) had approved all experiments and procedures. All experiments were approved and licensed by the relevant institutional committee, the Tierschutzbüro of the University Medical Center Göttingen (approval number T 09/08). Furthermore, methods are reported and carried out in accordance with ARRIVE guidelines and relevant regulations. Pubs were decapitated, brains were extracted and hippocampi dissected from both hemispheres. After removing the meninges, hippocampi were washed in 4°C cold Hank’s Balanced Salt Solution (HBSS, #14175-053, Invitrogen, Waltham, MA, USA), and afterwards incubated for 1 h at 37 °C in a slowly rotated (~30 rpm) enzyme solution, containing 15 U/ml papain (#LS003126, Worthington, Lakewood, USA), 0.5 mg/ml L-cysteine (#30090, Merck), 1 mM CaCl_2_ (#102382, Merck), and 0.5 mM EDTA (#108418, Merck) in DMEM, which was previously equilibrated with CO_2_ for 10 minutes. Inactivation of enzyme activity was achieved by transferring the hippocampi to a prewarmed enzyme inactivation solution, containing 5 mg/ml bovine serum albumin (BSA, #A1391, Applichem, Darmstadt, Germany) and 10% FCS in DMEM. After 15 minutes incubation at 37°C, the inactivating solution was replaced by pre-warmed plating medium (10% horse serum (#S900-500, VWR International GmbH, Darmstadt, Germany), 1.8 mM glutamine, 0.6 mg/ml glucose (#108342, Merck) in MEM (#51200046, ThermoFisher Scientific) and hippocampi were washed 3-4× with plating medium. Neurons were isolated by gentle trituration with a 10 ml serological pipette in 6 ml of plating medium, before they were centrifuged at 800 rpm for 8 minutes. Afterwards, supernatant was removed and cells were resuspended in 6 ml plating medium, before they were seeded at a density of 80k on PLL coated coverslips. Neurons could adhere to coverslips for 2-3 h in the incubator, then plating medium was replaced by Neurobasal culture medium (0.2% B27-supplement, #17504-044; 2 mM GlutaMAX #35050-038, in Neurobasal-A medium #10888-022, all ThermoFisher Scientific), and neurons were kept at 37°C and 5% CO2 for minimum 14 days, before they were fixed with 4% Paraformaldehyde (PFA #30525894, Merck) in PBS for 30 minutes at RT.

### Tubulin extraction and immunocytochemistry

Unfixed U2OS cells were incubated in a prewarmed solution of 0.2% Saponin (#47036, Sigma-Aldrich now Merck) in Cytoskeleton Buffer (CB, 10 mM 4-Morpholineethanesulfonic acid,2-(N-Morpholino)ethanesulfonic acid (MES #M3671, Merck), 138 mM KCl (#6781, Carl Roth, Karlsruhe, Germany), 3 mM MgCl_2_ (#105833, Merck), 2 mM EGTA (Triplex®VI #108435, Merck), 320 mM sucrose (#107651, Merck), pH:6.1) for 1 minute and were subsequently fixed with prewarmed 4% PFA and 0.1% Glutaraldehyde (#A3166, PanReac, Darmstadt, Germany) in CB. Quenching of unreactive aldehydes was achieved by 0.1% NaBH_4_ (#71320, Sigma-Aldrich now Merck) for 7 minutes, followed by incubation in 0.1 M glycine (#3187, Carl Roth) in PBS for 10 minutes. Immunocytochemistry was performed after premixing the primary anti-tubulin antibodies (Ab, #T6199 Sigma-Aldrich, #302211 Synaptic Systems, Göttingen, Germany, #302203 Synaptic Systems, and #ab18251 Abcam, Cambridge, UK), using AbberiorStar635P (AS635P) labeled secondary nanobodies (#N1202-Ab635P-S and #N2402-Ab635P-S both from NanoTag Biotechnologies, Göttingen, Germany) for 30 minutes at room temperature in a molar ratio of 1:5 (Ab:Nb). In the meanwhile, samples were blocked and permeabilized with 2% BSA and 0.1% Triton X-100 (#9036-19-5, Sigma-Aldrich now Merck) in PBS for 30 minutes at RT. The AB-Nb-mix was diluted in blocking/permeabilization buffer and was added to the cells for 1 h at room temperature. Samples were washed with PBS for 3× ten minutes and then further processed by expansion protocols.

### Vesicle plating

Synaptic vesicles were obtained from rat brains and isolated as described before [43,44]. For vesicle plating, clean and sterilized 18 mm glass coverslips were placed into a 12 well plate and coated with sterile-filtered 5% BSA in PBS at 37°C overnight. Afterwards, coverslips were washed 3x five minutes with PBS and 100-150 μl of vesicle solution was added to the coverslips in 500 μl PBS. The plate was centrifuged at 4000 rpm for 30 minutes at room temperature, coverslips were gently washed with PBS for five minutes, and then vesicles were fixed with 4% PFA in PBS for 30 minutes at room temperature. Afterwards, coverslips were quenched in NH4Cl (#101145100, Merck) for 20-30 minutes at RT and were used for immunocytochemistry.

### Tissue supply and immunohistochemistry of brain slices

Rat brains were obtained from P0-P1 old Wistar rat pups. All experiments were ap-proved by the local authority, the State of Lower Saxony (Landesamt für Ver-braucherschutz, LAVES, Braunschweig, Germany), as it is described in detail in section *Primary neuronal culture*. Rats were decapitated, brains were removed and directly fixed with 4% PFA for 20 h. Brains were embedded in 4% Agarose (#9012366, VWR Life Science, Hannover, Germany) and were cut into 100, 150 or 200 μm thick slices with a vibratome. After quenching with 50 mM glycine in PBS, slices were gently washed with PBS 3x for five minutes and blocked in 2.5% BSA in 0.3% Triton-PBS (PBS-T) for 2 h. Primary antibodies against Bassoon (#ADI-VAM-PS003-F, Enzo Life Sciences GmbH, Lörrach, Germany) and Homer1 (#160003, Synaptic Systems) were diluted 1:500 in 2.5% BSA in PBS and slices were incubated overnight at 4°C. After washing the slices for 3× five minutes in PBS, they were incubated in secondary antibodies AS635P (#ST635P-1001, Abberior, Göttingen, Germany), for the detection of Bassoon, and Cy3 (#711-165-152, Dianova, Hamburg, Germany) for the detection of Homer1, in a dilution of 1:1000 in 2.5% BSA in PBS-T for 3 h at room temperature. Finally, the slices were washed 5× for five minutes in 2.5% BSA in PBS-T and afterwards 2× for five minutes in PBS. Brain slices were then used for starting the expansion procedure.

### Immunocytochemistry of neurons and isolated vesicles

After fixation all neurons and isolated vesicles were quenched with 100 mM NH4Cl for 20 min at room temperature and blocked and permeabilized with 2.5 % BSA in PBS-T (PBS + 0.1 % Triton-X100, blocking/permeabilization solution, BSA-T) 3× for five minutes. Primary antibodies and nanobodies were diluted in BSA-T and samples were incubated in the anti-/nanobody solution for 1 h at room temperature. The primary antibodies used were anti synaptotagmin1 (SYT1, #105011 Synaptic Systems), anti synaptophysin1 (SYP1, #101004 Synaptic Systems), and anti SHANK2 (#162204 Synaptic Systems). Primary nanobodies were anti VGLUT1-biotinylated and anti SYT1 (#N1605-Biotin, #N2305-Biotin, both from NanoTag Biotechnologies, Göttingen). Anti VGLUT1-ALFA-tag and anti VGLUT1-AF488 (#N1605-ALFA, #N1605-AF488 were custom made by NanoTag Biotechnologies). After incubation with primary anti-/nanobodies, samples were washed 3× with BSA-T for 30 minutes at room temperature. The immunostainings could be combined with any of the amplification systems described later. Incubation in secondary anti-/nanobodies, without being amplified, was performed with either antibodies conjugated to Alexa Fluor 488 (AF488, #706-545-148, Dianova), Alexa Fluor 546 (AF546, #A11030, Invitrogen, Waltham, USA), Abberior Star 580 (AS580 #ST580-1006, Abberior) or a nanobody that was conjugated in house to Alexa Fluor 546 (see the following paragraph). After all anti-/nanobody incubations, samples were washed 3x for 10 minutes with PBS and were postfixed with 4% PFA in PBS for 15-20 minutes at room temperature. Afterwards, samples were again quenched with 100 mM NH4Cl for 10 minutes at room temperature and kept in PBS until further processing.

### Nanobody conjugation to Alexa Fluor546

Unconjugated single-domain antibody anti-mouse IgG1 was obtained from NanoTag Biotechnologies (#N2005). The nanobody was reduced using 10 mM Tris(2-carboxyethyl)phosphin -hydrochlorid (TCEP, #51805-45-9, Merck) on ice for 1h. The excess of TCEP was removed using a NAP5 column (Cytiva, Washington, USA) equilibrated with ice-cold PBS buffer, pH7.4, and mixed immediately with 3 molar excess of maleimide Alexa Fluor546 (#A10258, Thermo Scientific), followed by incubation at room temperature for another 2h. The conjugated nanobody was separated from free fluorophore using a Superdex 75 Increase 10/300 GL column (Cytiva) equilibrated with 2× PBS pH7.4. Conjugated fractions were pooled, the concentration was measured and the sample was diluted with glycerol (#2039, Chemsolute by Th. Greyer, Renningen, Germany) to reach 50% glycerol in 1× PBS, enabling the long-term storage in aliquots at −80°C.

### Application of the AF488 amplification system

Samples containing labeling with AF488 could be further incubated with an antibody against AF488 (#A-11094, ThermoFisher Scientific) in BSA-T for 30 minutes at room temperature. A 3× wash with BSA-T was performed before samples were incubated in the final secondary antibody in BSA-T for 30 minutes at room temperature. Antibodies used here were either AF488 (#711-545-152, Dianova) or AS635P (#ST635P-1002, Abberior). Afterwards, samples were washed with PBS and postfixed, as described above.

### Biotin amplification system

The immunostaining with primary nanobodies conjugated to biotin (BT) was performed as described above, relying on 30 minutes of incubation at room temperature with an antibody against BT (#31852, ThermoFisher Scientific), diluted in BSA-T. Afterwards, samples were washed 3x for five minutes with BSA-T, before the incubation with the last antibody carrying AS635P (#ST635P-1055, Abberior), AS580 (#ST580-1002, Abberior) or AF488 (#705-545-147, Dianova). This antibody was applied for 30 minutes at room temperature. Final washes with PBS and postfixation were performed as described above.

### ALFA and Spaghetti Monster amplification system

After immunostaining the samples with primary nanobodies carrying an ALFA-tag, two possible procedures were followed to reveal the ALFA-tag in our X10ht methodology. For the condition termed “BEFORE AC”, the samples were incubated for 30 minutes at room temperature with a NbALFA (#N1502, NanoTag Biotechnologies) as it is performed for general immunostainings directly after a 3× wash with BSA-T. The NbALFA was produced fused to a protein called Spaghetti Monster (SpaMo, Viswanathan et al., 2015), which is an engineered GFP containing 7 FLAG-tags (custom made by NanoTag Biotechnologies). Following the incubation with the SpaMo, samples were washed 3× five minutes in BSA-T and incubated with an antibody against the FLAG-tag (#F1804 Sigma-Aldrich, now Merck, or #14793 Cell signaling, Leiden, the Netherlands), for 30 minutes at room temperature. After repeated washing with BSA-T, samples were incubated with fluorophore-conjugated secondary antibodies (AS635P #ST635P-1001; AS580 #ST580-1001, both Abberior; Cy3 #715-165-150, Dianova) or nanobodies (AS635P, #N1202-Ab635P, NanoTag Biotechnologies) for 30 minutes at room temperature, before washing with PBS and post-fixation, as described above.

Alternatively, we labelled the ALFA-tagged nanobodies during the expansion procedure (for better understanding see scheme in Fig. 5A). For the “AFTER AC” immunostaining, only the primary nanobody carrying the ALFA-tag was applied during the general immunocytochemistry (pre-expansion). The samples underwent the expansion procedure (which is described in detail below), stopped after the samples were autoclaved (AC). At this time point, the samples were incubated with the NbALFA-SpaMo in BSA-T overnight at room temperature on a slowly rotating shaker. The next day, samples were washed carefully in PBS (5×, 1-2 hours) and incubated in anti-FLAG-tag antibody in BSA-T overnight at room temperature on a shaker. Unbound antibody was washed away with PBS 5× for 1-2 h, before the last antibody immunostaining was performed, with fluorophore-conjugated secondary anti- or nanobodies. This final step was performed in BSA-T overnight at room temperature, on a slow rotating shaker. Alternatively, using the secondary AS635P nanobody, a premixing of the anti-FLAG-tag antibody with the respective nanobody (30 minutes at room temperature in a 1:5 molar ratio antibody:nanobody), which helps to reduce the incubation step to one overnight incubation, instead of two. The next day, samples were ready to continue with the last steps of the expansion procedure (see later paragraphs).

### Expansion procedure

Immunostained samples were processed as described in Truckenbrodt et al. 2019, with some alterations of the protocol, which is termed X10ht in the following paragraphs [8]. For maximal anchoring of the proteins and labels to the gel, we used 0.3 mg/ml Acryloyl-X (SE; #A-20770, ThermoFisher Scientific) in 150 mM NaHCO_3_ buffer (Merck, #144-55-8) with a pH of 11. Anchoring was performed overnight on a shaker at room temperature. Over the next day samples were washed 3× with PBS for five min, while the monomer solution was prepared as described before [25]. Briefly, 1.335 g *N,N*-dimethylacrylamide (DMAA, #274135, Sigma-Aldrich now Merck) and 0.32 g sodium acrylate (SA, #408220, Sigma-Aldrich now Merck) were dissolved in 2.850 g ddH_2_O and purged with N2 for 40 minutes. Afterwards, 300 μl of a 0.36 g/ml potassium persulfate (KPS, #379824, Sigma-Aldrich now Merck) stock solution was added to 2700 μl of the DMAA/SA solution and further purged with N_2_ on ice for 15 minutes. Immediately before usage, 4 μl of *N,N,N’,N’*-Tetramethylethylendiamine (TEMED, #612-103-00-3, Sigma-Aldrich now Merck) was added to 1 ml of the monomer solution, was vortexed and drops were pipetted on parafilm, on which the specimen was placed. Coverslips with cells were flipped to incubate cells in the solution, while drops were applied directly on the tissue slices and covered with a coverslip. Gels polymerized for 6-20 h at a stable temperature of 23°C in a humidified chamber.

The homogenization with 8 U/ml proteinase K (PK, #P4850 Sigma-Aldrich now Merck) in digestion buffer (50 mM TRIS (#AE15.2, Carl Roth GmbH), 800 mM guanidine HCl (#G3272, Sigma-Aldrich now Merck), 2 mM CaCl_2_ and 0.5 % Triton X-100) was performed as described before [8,25]. This procedure was only performed for the gels used for images labeled “PK” in our Figures, and was not combined with heating-based homogenization.

In the present study, we present a heat homogenization step for 10X expansion by the application of a milder protein disruption via autoclaving over 100°C (AC), naming the new protocol 10Xht. Polymerized gels were rinsed shortly with 1 M NaCl (#7647145, Merck) and afterwards extensively soaked in disruption buffer, consisting of 100 mM TRIS, 5% Triton-X and 1% SDS (#1057.1, Carl Roth GmbH) in ddH_2_O with pH 8, for minimal 2 h at room temperature, while buffer was exchanged 4×. Autoclaving was performed for 30 minutes at 110-121°C and gels were allowed to cool down slowly. To enable the overview imaging of the expanded tissue, slices were labeled with NHS-Fluorescein (#46409, ThermoFisher Scientific) over night after being extensively washed with PBS for 5× for 30 minutes. For expanding the gels after homogenization (both PK-digested and autoclaved gels) or NHS-Fluorescein, they were placed in 22 × 22 cm culture dishes and ~ 400 ml ddH_2_O was added to each gel. Water was exchanged every 30 – 90 minutes for 4× in total and gels were left in water over night to fully expand.

### Investigation of optimal temperature for X10ht

Fixed BHK cells were processed for expansion as described in Truckenbrodt et al., 2018 until homogenization. Instead of applying proteinase K in digestion buffer, gels were shortly rinsed with 1 M NaCl and soaked in disruption buffer for ~ 2h while exchanging the buffer 4x. Afterwards, gels were autoclaved for 30 minutes at either 70, 80, 90, 100, or 110°C. Disruption buffer needed to be extensively washed away before adding 1 ml 1 μM Atto590-NHS ester (NHS-Atto590, #79636, Merck) in PBS to each gel, to label all retained proteins, followed by incubation overnight at room temperature on a slowly rotating shaker. On the next day, gels were expanded as it was described before [8] and imaged.

### Image acquisition

For image acquisition, gels were cut into smaller pieces and placed in a self-made imaging chamber on top of a glass coverslip. Water was removed to the maximum by simply soaking any unnecessary droplets with paper tissues. Most epifluorescence images were obtained with an inverted epifluorescence microscope, Nikon Eclipse Ti (Nikon Corporation), using a Plan Apochromat 60x objective (1.4 NA, oil immersion), a HBO-100W Lamp, and relying on an IXON X3897 Andor camera. Alternatively, we used a second inverted fluorescence microscope, Olympus IX 71 (Olympus, Hamburg, Germany), using a 20× objective (from Olympus). The Olympus setup employed a CCD camera (FView II, Olympus) and CellF software (Olympus). To identify NHS-AF590, a 545/30 HQ excitation filter (AHF, Tübingen, Germany), a 570 LP Q beam splitter and a 610/75 HQ emission filter were used.

An Abberior Expert line setup (Abberior Instruments) using an IX83 microscope (Olympus) with an 100× oil immersion objective (UPLSAPO, 1.4 NA; Olympus) was employed to generate confocal and STED images. Star635P was excited with a 640 nm excitation laser (set to 10-30% of max. power, nominally 1.77 mW, pulsed at 80 MHz). Signals were detected using an avalanche photodiode (APD) that has a pre-set range of 650-720 nm. A 561 nm excitation laser (40-50% of max. power, 440 μW; 80 MHz) was used to excite AS580 or Cy3, while the emission was detected with an APD, 605-625 nm. Depletion for AS635P, AS580 or Cy3 was achieved by a 775 nm depletion laser (set to 15-40% of max. power of 1.2 W). Excitation of AF488 was achieved by a 485 nm excitation laser (10-15% of max. power), and detected with an APD 525-575 nm. The used depletion laser for AF488 was a pulsed solid-state 595 nm laser (set to ~20% of max. power of 2 W). For all images, the pixel size was 20 nm, with a dwell time of 5-10 μs per pixel and a line accumulation of 3-5 for STED.

### Immunocytochemistry, expansion and analysis of Nup96

In order to visualize expanded nuclear pore complexes, U2OS cells expressing mEGFP coupled to NUP96 were fixed, permeabilized and blocked as described above. Cells were incubated in a mixture of two different anti-GFP nanobodies, which were conjugated to Alexa Fluor 488 (#N0302-AF488, custom made; #N0303-AF488, custom made, both NanoTag Biotechnologies) for 1h at room temperature. Afterwards, they were washed 3× for five minutes with blocking solution and the AF488 amplification immunostaining was performed as described above. The last secondary antibody was conjugated to AS635P (#ST635P-1002, Abberior). Cells were washed 3× for five minutes with PBS and post-fixed before the expansion microscopy X10ht protocol was applied, followed by imaging with the Abberior STED setup. STED images were analyzed with a custom-made Matlab script, which measured the distances between all fluorescent localizations. Plot profiles of exemplary line scans were generated with ImageJ and data are plotted using GraphPad Prism 7.

### Analysis of X10ht expansion factor of U2OS nuclei diameter

Fixed U2OS cells were treated by following either the original X10 protocol with PK digestion [8] or the X10ht protocol with the usage of heat denaturation (AC). After the respective homogenization procedure, gels were washed for around 2-4 h with PBS and incubated in 1 ml 1 μM Alexa Fluor 546 NHS ester (NHS-AF546, #A20002, Merck), overnight at room temperature on a slowly rotating shaker. The next day expansion of gels was performed like described before. For image acquisition, the Nikon Eclipse Ti epifluorescence microscope was used as described above. The measurements of nucleus diameter was performed by drawing vertical lines from the highest to the lowest point of each nucleus, using ImageJ. As control, the same measurements were performed in non-expanded cultures (-Ex). For this procedure, fixed cells were incubated in Hoechst (1:2000 in PBS, #62249, Thermo Scientific) for 5 minutes at room temperature, washed 3× for 10 minutes in PBS and embedded in Mowiol (#0713.2, Carl Roth GmbH).

### Image processing, analysis and statistics

In case of drifting, images were manually drift-corrected with ImageJ (Wayne Rasband, NIH, Bethesda, MD, USA). Signal intensities of images obtained by epifluorescence microscopy were analyzed with ImageJ by shrinking the average bin size of the raw file and measuring the total grey values of 5-30 pictures per condition. Fluorescence intensity evaluation of confocal or STED images was performed on raw image files with a semi-automated custom-written Matlab (The Mathworks, Inc, Natick, MA, USA) routine, analyzing mean grey values per ROI. Line scans and plot profiles of the images presented in Fig. 2 were performed using Matlab.

Spot resolution was analyzed by performing line scans of 0.6 μm on fluorescent events with Matlab (Fig. 6). FWHM was calculated by plotting the line scans and employing a Lorentzian fit. Distance analysis of vesicular fluorescent spots were performed with ImageJ.

All scatter plots were generated and statistics were performed using GraphPad Prism 7 (GraphPad Software Inc., La Jolla, CA, USA). Data were analyzed for normal distribution by the Shapiro Wilk-test and were then tested for significance with students-t test (when normally distributed) or the Mann Whitney-U-test (when not normally distributed). Data is presented as individual data points and means ± SD. Differences were considered significant and were indicated by asterisks with *P<0.05; **P<0.01; ***P<0.001, and ****P<0.0001. Illustrations for Fig. 3 were created with BioRender (BioRender.com; Science Suite Inc., Canada) and figures were prepared with Adobe Illustrator CS6 (Adobe Systems Incorporated, Mountain View, USA).

