## supplementary for "Heat denaturation enables multicolor X10-STED microscopy at single-digit nanometer resolution"

### Table of content

#### Supplementary Figures


- **Supplementary Fig. A1:** Optimization of autoclaving temperature

- **Supplementary Fig. A2:** Comparison of expansion factor after the enzymatic digestion and autoclaving methods

- **Supplementary Fig A3:** Application of the X10ht protocol applied to the investigation of Nup96 in U2OS-mEGFP cells

- **Supplementary Fig. A4:** Exemplary confocal images show the application of different signal amplification systems in primary neuronal cultures, using different fluorophores

- **Supplementary Fig. A5:** Analysis of 3-color STED images to identify isolated synaptic vesicles in X10ht

**Supplementary figures**

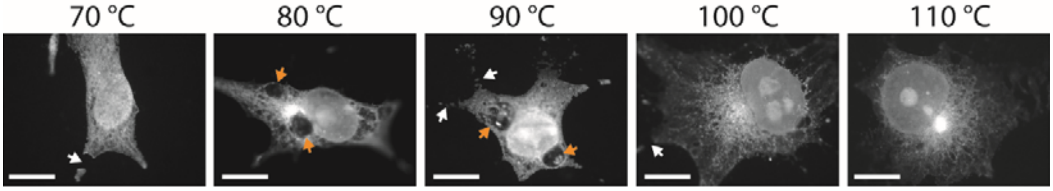


***Figure A1: Optimization of autoclaving temperature.*** *Exemplary epifluorescence images of expanded BHK cells are shown, treated with ascending temperatures from 70 – 110 °C during autoclaving. Lower temperatures resulted in cracked cells (white arrows), large cytosolic swellings (orange arrows), while higher temperature ensured the integrity of the cells and a higher expansion size. All cells are stained with NHS-Atto590 after autoclaving. Scale bar 100 µm, without taking into consideration the expansion factor.*

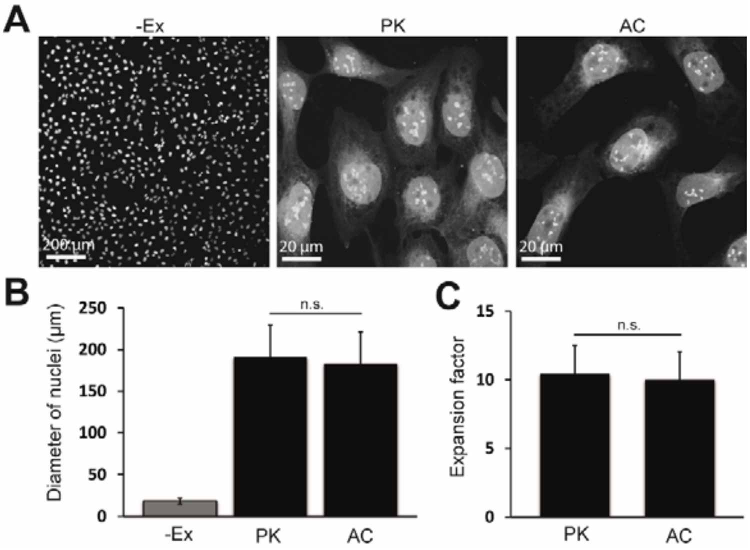


***Figure A2: Comparison of expansion factor after the enzymatic digestion and autoclaving methods. A)*** *Exemplary epifluorescence images for not expanded U2OS cells (-Ex) and after expansion with enzymatic homogenization with proteinase K (PK) or autoclaving at 110 °C (AC). All cells are stained with NHS-AF546 after homogenization.* ***B)*** *The analysis of the nucleus size revealed no significant (n.s.) difference in the size of expansion between the two protocols, and an expansion factor of ~10 for both conditions (shown in* ***C****). N = 29 analyzed cells for –Ex, 74 analyzed cells for PK and 49 analyzed cells for AC obtained from two independent experiments.*

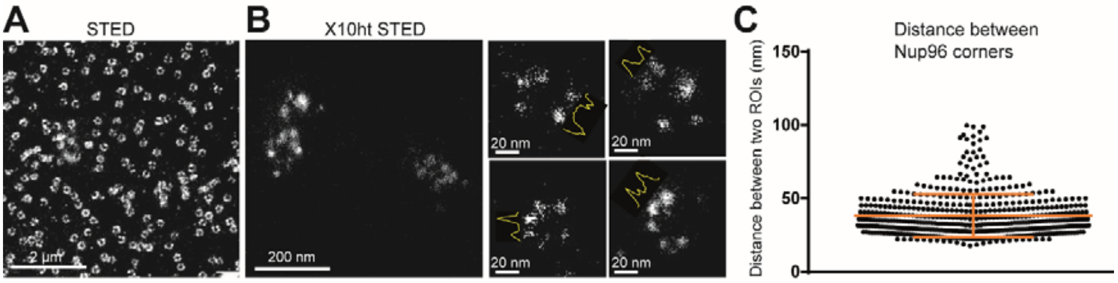


***Figure A3: Application of the 10Xht protocol applied to the investigation of Nup96 in U2OS-mEGFP cells. A)*** *STED image of not expanded Nup96 (STED).* ***B)*** *Nup96 ring-like structures after X10ht, imaged in STED. Additional examples are shown in the smaller panels, with the line scans in yellow depicting the profiles of two adjacent corners of a NPC.* ***C)*** *Quantification of the distance between the corners (mean value ~38 nm). N = 516 Nup96 subunits from 58 images were analyzed.*

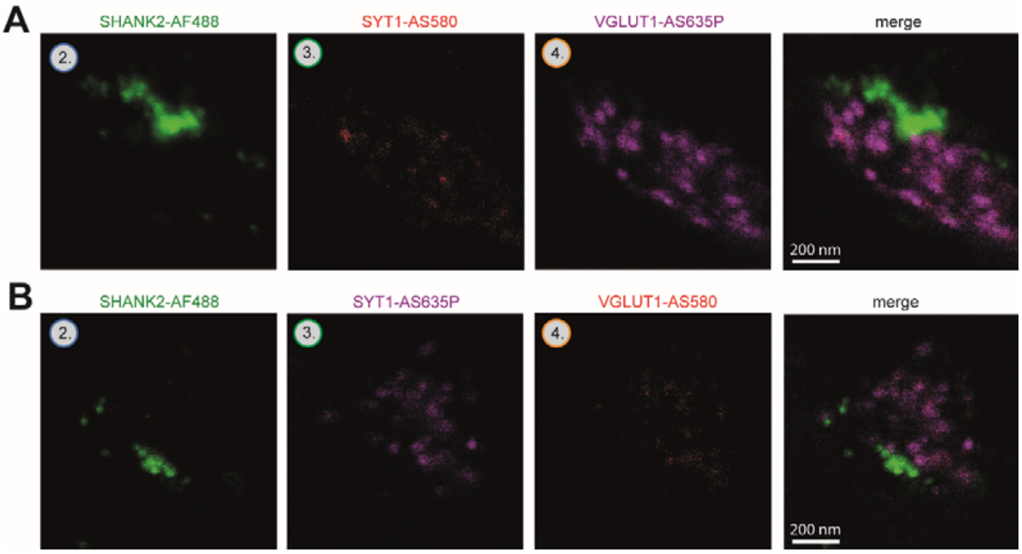


***Figure A4: Exemplary confocal images show the application of different signal amplification systems in primary neuronal cultures, using different fluorophores. A)*** *The combination of the AF488 system was used for labeling postsynaptic compartments (SHANK2) in green, together with NbSYT1-BT + AbAS580 and the ALFA-SpaMo amplification to depict VGLUT1 with AbAS635P.* ***B)*** *The same immunostaining with a different combination of fluorophores for SYT1 and VGLUT1.*

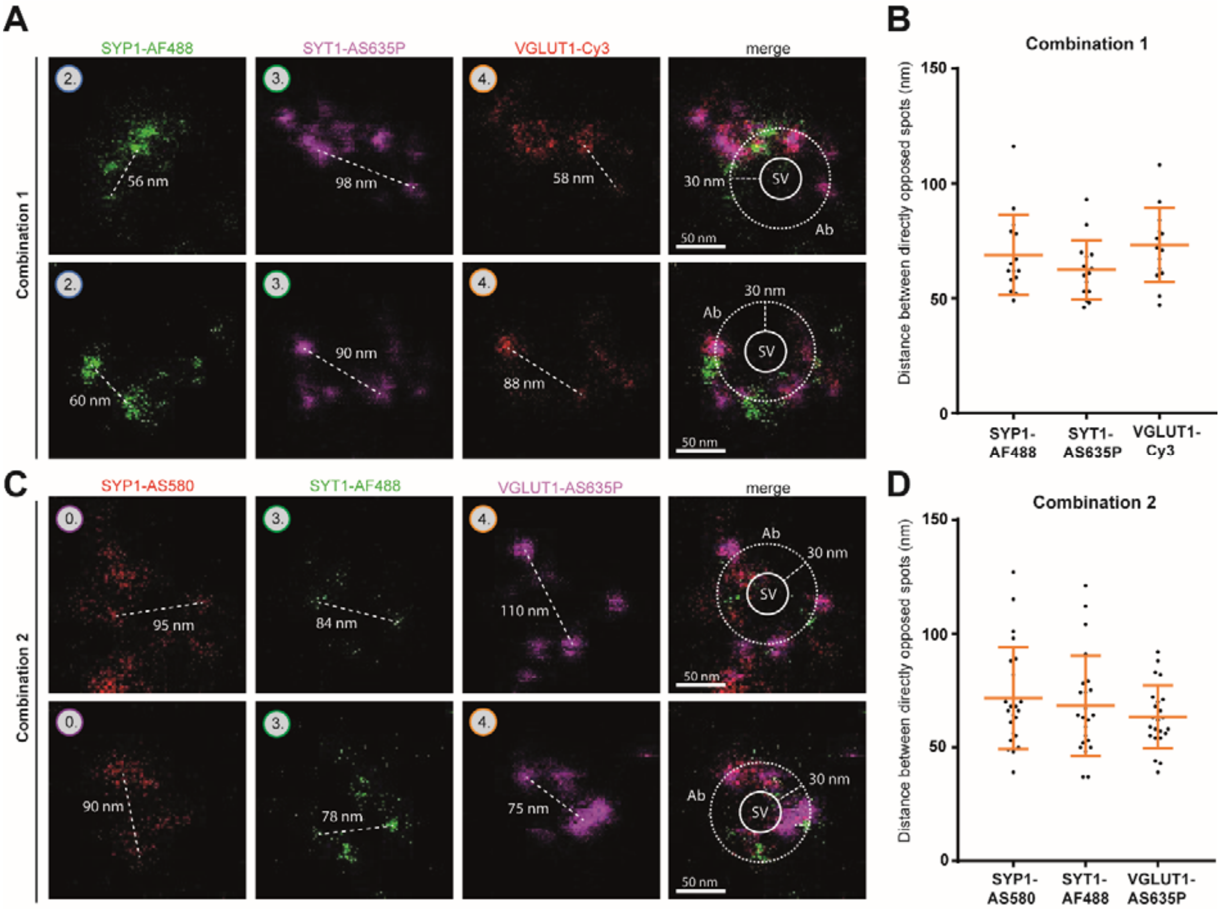


***Figure A5: Analysis of 3-color STED images to identify isolated synaptic vesicles in X10ht. A)*** *Two representative examples of the vesicle proteins SYP1, SYT1 and VGLUT1, immunostained with different amplification systems and fluorophore combinations.* ***B)*** *An analysis of distances between the spots (white dashed lines in A) of each channel. N = 15 images with several spot distances were analyzed.* ***C)*** *The same analysis, for a different fluorophore combination, with a similar quantification in panel* ***D****. N = 21 images with several spot distances were analyzed. In the merged pictures, the synaptic vesicle (SV) is depicted by the inner circle (continuous line) and the region where the antibodies (Ab) are probably located is illustrated as the larger dashed circle. Data are presented given as individual data points, mean ± SD.*
